## Supplementary material for "A transthalamic pathway crucial for perception": Mo et al_Supplement

**Supplemental Material**

**Extended Methods**

Anatomical Verification of the S1 L5 to Pom to S2 Pathway

With regards to **Suppl. Fig. 1**, Rbp4-Cre mice were injected with Cre-dependent Jaws-TdTomato and implanted with a head bar and cranial window above S1 and S2, as described in **Surgical procedures**. IS imaging was performed to identify S1 and S2 (see **Intrinsic signal optical imaging**) and 80nl of the retrograde tracer, FluoroGold, was injected into S2. One week later, the mice were perfused and sectioned (50µm thick) to locate the region of POm with overlapping Jaws-TdTomato and Fluorogold cell body expression.

In Vivo Electrophysiology

With regards to **Suppl. Fig. 2**, we validated Jaws opsin inhibition at S1 L5 to POm terminals. Jaws and ChR2 was expressed in S1 L5 neurons by injecting 450nl of a 1:1 mix of AAV8-CAG-FLEX-Jaws-KGC-TdTomato-ER2 and AAV5-DIO-ChR2-eYFP (UNC Vector Core) in Rbp4-Cre mice (n=4). After 3 - 4 weeks, mice were anesthetized with urethane (1.3mg/g) and a 6mm x 5mm craniotomy was made over S1 and POm. A 473nm LED was placed over S1 (ML: 3.1mm, AP: -0.8) and multiunit recordings were made in POm (DV: −3.1, ML: +1.3, AP: −1.4 mm) using custom pulled tungsten-in-glass microelectrodes (1Mohm). The microelectrode was attached to an optic fiber implant (200μm diameter, 0.5NA, Thorlabs) such that the tip of the electrode was within 290μm of the end of the implant. The implant was connected to a 620nm LED (max 137mW/mm^2^, PlexBright, Plexon). Custom Matlab code coordinated 473nm LED pulses (1Hz, 20msec long) and 620nm LED pulses (1-3sec long) with various relative onset times (50msec, 500msec, 1sec, 2sec). Signals were passed through a 300Hz high-pass and 12000Hz low-pass second-order Butterworth filter. The window for spike analysis was 6-40msec from the start of the 473nm pulse and spikes were thresholded to 5 standard deviations above baseline.

**Supplemental table 1. Statistical analysis of data**

| **Figure** | **Data (mean ± sem)** | **P-value** | **Statistical test** |
| --- | --- | --- | --- |
| **Fig. 2B** | **Error rate No-laser vs Sensory laser**  0.23 ± 0.013 vs 0.41 ± 0.023 | p=0.0039 | Paired samples Wilcoxon test |
| **Fig. 2C** | **Psychometric curve No-laser vs Sensory laser**  effect of texture  effect of laser  texture x laser interaction  *No-laser vs Sensory laser*  G0: 26.54 ± 2.14 vs 44.44 ± 3.67  G1: 43.29 ± 2.60 vs 45.64 ± 3.28  G2: 54.67 ± 3.69 vs 53.78 ± 3.03  G3: 74.44 ± 3.68 vs 64.14 ± 5.58  G4: 74.16 ± 3.56 vs 60.73 ± 4.84  G5: 79.17 ± 2.52 vs 63.20 ± 4.35 | p<0.0001  p=0.094  p<0.0001  p=0.0078  p>0.99  p>0.99  p=0.18  p=0.017  p=0.0016 | 2-way RM ANOVA  Bonferroni post-hoc tests |
|  | **Psychometric parameters No-laser vs Sensory laser**  lapse rate: 0.22 ± 0.031 vs 0.37 ± 0.044  guess rate: 0.20 ± 0.041 vs 0.40 ± 0.039  bias: 2.51 ± 0.23 vs 2.49 ± 0.33  sensitivity: 1.11 ± 0.20 vs 0.11 ± 0.10 | p=0.0039  p=0.0078  p=0.82  p=0.0039 | Paired samples Wilcoxon test |
| **Fig. 2C** | **D-prime No-laser vs Sensory laser**  effect of texture  effect of laser  texture x laser interaction  *No-laser vs Sensory laser*  G1: 0.47 ± 0.035 vs 0.029 ± 0.072  G2: 0.76 ± 0.085 vs 0.24 ± 0.076  G3: 1.33 ± 0.10 vs 0.54 ± 0.17  G4: 1.32 ± 0.12 vs 0.43 ± 0.15  G5: 1.48 ± 0.087 vs 0.50 ± 0.12 | p<0.0001  p<0.0001  p=0.0007  p=0.0012  p=0.0065  p=0.0012  p=0.0015  p<0.0001 | Two-way repeated measures (RM) ANOVA  Bonferroni post-hoc tests |
| **Fig. 2D** | **D-prime performance across trial conditions**  effect of condition  No-laser vs Sensory laser: 1.43 ± 0.080 vs 0.50 ± 0.12  No-laser v Delay laser: 1.43 ± 0.080 vs 0.95 ± 0.13  No-laser vs No whiskers: 1.43 ± 0.080 vs 0.28 ± 0.071 | p<0.0001  p<0.0001  p=0.025  p=0.025 | Mixed-effects model  Bonferroni post-hoc tests |
| **Fig. 2E** | **Error rate No-laser vs Delay laser**  0.25 ± 0.017 vs 0.32 ± 0.023 | p=0.0055 | Paired samples Wilcoxon test |
| **Fig. 2F** | **Psychometric curve No-laser vs Delay laser**  effect of texture  effect of laser  texture x laser interaction  *No-laser vs Delay laser*  G0: 29.14 ± 1.91 vs 35.06 ± 2.41  G1: 41.04 ± 1.80 vs 38.69 ± 2.84  G2: 62.26 ± 2.36 vs 50.92 ± 3.14  G3: 71.58 ± 2.25 vs 66.84 ± 3.72  G4: 73.84 ± 1.70 vs 70.09 ± 2.80  G5: 78.82 ± 2.11 vs 70.59 ± 3.18 | p<0.0001  p=0.0098  p=0.027  p=0.055  p>0.99  p=0.23  p=0.99  p>0.99  p=0.35 | 2-way RM ANOVA  Bonferroni post-hoc tests |
|  | **Psychometric parameters No-laser vs Delay laser**  lapse rate: 0.24 ± 0.016 vs 0.30 ± 0.027  guess rate: 0.30 ± 0.032 vs 0.37 ± 0.030  bias: 2.73 ± 0.13 vs 3.35 ± 0.14  sensitivity: 0.057 ± 0.29 vs 0.0036 ± 0.13 | p=0.027  p=0.016  p=0.039  p=0.91 | Paired samples Wilcoxon test |
| **Fig. 2F** | **D-prime No-laser vs Delay laser**  effect of texture  effect of laser  texture x laser interaction  *No-laser vs Delay laser*  G1: 0.33 ± 0.056 vs 0.097 ± 0.063  G2: 0.87 ± 0.083 vs 0.41 ± 0.11  G3: 1.14 ± 0.092 vs 0.85 ± 0.14  G4: 1.2 ± 0.084 vs 0.93 ± 0.10  G5: 1.38 ± 0.11 vs 0.95 ± 0.13 | p<0.0001  p=0.0011  p=0.36  p=0.023  p=0.048  p=0.10  p=0.029  p=0.11 | Two-way RM ANOVA  Bonferroni post-hoc tests |
| **Fig. 2G** | **POm implant No-laser vs Sensory vs Delay laser**  effect of texture  effect of laser  texture x laser interaction  *No-laser vs Sensory laser*  G0: 0.30 ± 0.024 vs 0.43 ± 0.027  G1: 0.45 ± 0.029 vs 0.49 ± 0.026  G2: 0.70 ± 0.034 vs 0.60 ± 0.039  G5: 0.80 ± 0.015 vs 0.66 ± 0.041  *No-laser vs Delay laser*  G0: 0.30 ± 0.024 vs 0.38 ± 0.029  G1: 0.45 ± 0.029 vs 0.46 ± 0.023  G2: 0.70 ± 0.034 vs 0.55 ± 0.029  G5: 0.80 ± 0.015 vs 0.72 ± 0.030  **Control implant**  effect of texture  effect of laser  texture x laser interaction  *No-laser vs Sensory laser*  G0: 0.34 ± 0.042 vs 0.40 ± 0.023  G1: 0.43 ± 0.044 vs 0.44 ± 0.032  G2: 0.67 ± 0.050 vs 0.67 ± 0.039  G5: 0.76 ± 0.026 vs 0.74 ± 0.035  No-laser vs Delay laser  G0: 0.34 ± 0.042 vs 0.31 ± 0.064  G1: 0.43 ± 0.044 vs 0.46 ± 0.053  G2: 0.67 ± 0.050 vs 0.60 ± 0.060  G5: 0.76 ± 0.026 vs 0.72 ± 0.057 | p<0.0001  p=0.12  p<0.0001  p=0.0007  p=0.90  p=0.15  p=0.0086  p=0.14  p>0.99  p=0.0047  p=0.040  p=0.0027  p=0.16  p=0.43  p=0.91  p<0.999  p>0.999  p=0.72  p>0.999  p>0.999  p=0.26  p>0.999 | Two-way RM ANOVA  Bonferroni post-hoc tests  Two-way RM ANOVA  Bonferroni post-hoc tests |
| **Fig. 2H** | **Jaws and behavior correlation**  r: 0.73  R-squared: 0.51 | p=0.030  p=0.030 | Pearson’s correlation  Simple linear regression |
| **Fig. 2I** | **Whisking**  effect of trial  effect of laser  trial x laser interaction | p=0.056  p=0.63  p=0.79 | Two-way ANOVA |
| **Fig. 3D** | **Detection parameters Jaws No-laser vs Laser**  lapse rate: 0.17 ± 0.044 vs 0.20 ± 0.039  guess rate: 0.28 ± 0.47 vs 0.28 ± 0.047  bias: 4.21 ± 0.44 vs 5.02 ± 0.28  sensitivity: 1.10 ± 0.58 vs 0.69 ± 0.27 | p=0.22  p=0.91  p=0.048  p=0.49 | Paired t-test |
| **Fig. 3F** | **Detection parameters No-Jaws No-laser vs laser**  lapse rate: 0.21 ± 0.072 vs 0.19 ± 0.028  guess rate: 0.27 ± 0.021 vs 0.28 ± 0.026  bias: 4.20 ± 0.32 vs 4.50 ± 0.15  sensitivity: 0.88 ± 0.39 vs 0.91 ± 0.26 | p=0.83  p=0.77  p=0.47  p=0.96 | Paired t-test |
| **Fig. 4A** | **Behavioral performance of 2-photon imaging mice**  effect of texture  effect of sensory laser  texture x laser interaction  *No-laser vs Sensory laser*  G0: 0.23 ± 0.03 vs 0.43 ± 0.048  G1: 0.45 ± 0.048 vs 0.51 ± 0.052  G2: 0.58 ± 0.054 vs 0.51 ± 0.054  G5: 0.70 ± 0.02 vs 0.56 ±0.025 | p<0.0001  p=0.6958  p=0.0001  p=0.0071  p>0.099  p=0.76  p=0.0002 | Two-way RM ANOVA  Bonferroni post-hoc tests |
|  | **Overall texture responsiveness**  effect of brain region  effect of laser  brain region x laser interaction | p=0.51  p=0.11  p=0.27 | Two-way RM ANOVA |
| **Fig. 4C** | **Texture responsiveness: hit vs CR** *S1* effect of laser effect of texture laser x texture interaction *S2* effect of laser effect of texture laser x texture interaction No-laser: G5 Hit vs. No-laser:G0 CR No-laser: G5 Hit vs. Laser:G5 Hit No-laser: G5 Hit vs. Laser:G0 CR No-laser: G0 CR vs. Laser:G5 Hit No-laser: G0 CR vs. Laser:G0 CR Laser: G5 Hit vs. Laser:G0 CR | p=0.56  p=0.0033  p=0.91  p=0.64  p=0.31  p=0.0002  p=0.0013  p=0.032  p>0.99  p>0.99  p=0.68  p=0.86 | Two-way RM ANOVA  Bonferroni post-hoc tests |
| **Fig. 4D** | **Scatterplots of hit vs CR responsiveness**  *S1 no-laser vs laser*  No-laser r=0.69  Laser r=0.65  No-laser R-squared: 0.48  Laser R-squared: 0.43  S1 Slopes different  *S2 no-laser vs laser*  No-laser r=0.72  Laser r=0.77  No-laser R-squared: 0.52  Laser R-squared: 0.59  S2 Slopes different  **Scatterplots of hit vs FA responsiveness**  *S1 no-laser vs laser*  No-laser r=0.63  Laser r=0.69  No-laser R-squared: 0.39  Laser R-squared: 0.48  S1 Slopes different  *S2 no-laser vs laser*  No-laser r=0.51  Laser r=0.61  No-laser R-squared: 0.26  Laser R-squared: 0.37  S2 Slopes different | p<0.0001  p<0.0001  p<0.0001  p<0.0001  p=0.24  p<0.0001  p<0.0001  p<0.0001  p<0.0001  p<0.0001  p<0.0001  p<0.0001  p<0.0001  p<0.0001  p=0.59  p<0.0001  p<0.0001  p<0.0001  p<0.0001  p=0.0002 | Pearson’s correlation  Simple linear regression  Pearson’s correlation  Simple linear regression  Pearson’s correlation  Simple linear regression  Pearson’s correlation  Simple linear regression |
| **Fig. 4F** | **Fraction positive or negative DI cells**  *S1*  no-laser G5: 0.064 ± 0.016 vs no-laser G0: 0.019 ± 0.0090  laser G5: 0.03 ± 0.012 vs laser G0: 0.034 ± 0.018  no-laser G5: 0.064 ± 0.016 vs laser G5: 0.03 ± 0.012  *S2*  no-laser G5: 0.084 ± 0.017 vs no-laser G0: 0 ± 0  laser G5: 0.021 ± 0.014 vs laser G0: 0.072 ± 0.024  no-laser G5: 0.084 ± 0.017 vs laser G5: 0.021 ± 0.014  no-laser G0: 0 ± 0 vs laser G0: 0.072 ± 0.024 | p=0.016  p=0.27  p=0.051^  p=0.0009  p=0.18  p=0.025^  p=0.0013^ | Paired mutually exclusive McNemar’s test  ^ z-test |
|  | **Total fraction selective DI cells**  *No-laser vs laser*  S1: 0.082 ± 0.018 vs 0.063 ± 0.038  S2: 0.084 ± 0.017 vs 0.080 ± 0.022 | p=0.62  p=0.87 | Wilcoxon matched-pairs signed rank test |

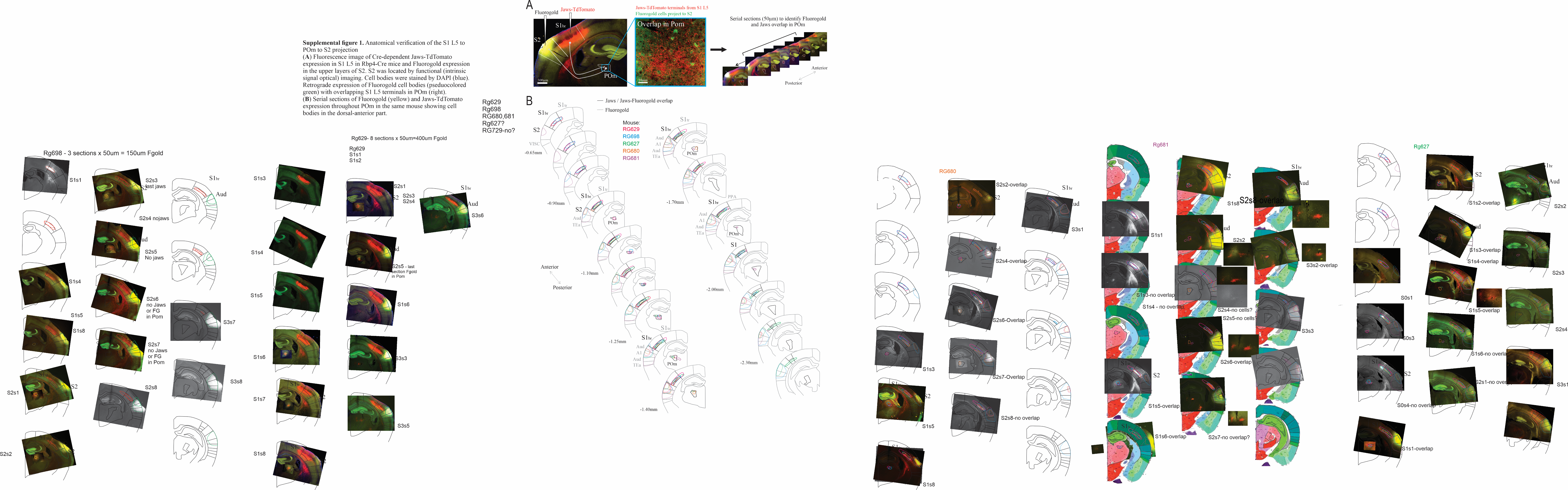

**Supplemental figure 1.** **Anatomical verification of the S1 L5 to POm to S2 projection**

(**A**) (Left) Injection strategy to label the feedforward transthalamic pathway: Jaws-TdTomato injected in S1 L5 and retrograde tracer Fluorogold injected in the upper layers of S2 in Rbp4-Cre mice. S2 was located by functional (intrinsic signal optical) imaging. (Middle) Retrograde expression of Fluorogold cell bodies (pseduocolored green) with overlapping S1 L5 terminals in POm. (Right) Sectioning to identify Jaws and Fluorogold overlay in POm.

(**B**) Coronal sections of Jaws-TdTomato and Fluorogold (dotted outlines) expression. Each color represents one experiment per mouse. POm is outlined in thalamus. Note the overlapping area in POm is in the anterior half of the nucleus. Approximate anterior-posterior coordinates relative to Bregma are indicated.

Aud: auditory area, A1: primary auditory cortex, POm: posterior medial nucleus, PPA: posterior parietal association areas, S1: primary somatosensory cortex, S1tr: trunk area of S1, S1br: barrel field (whisker area) of S1, S2: secondary somatosensory cortex, TEa: Temporal association areas, VISC: Visceral area.

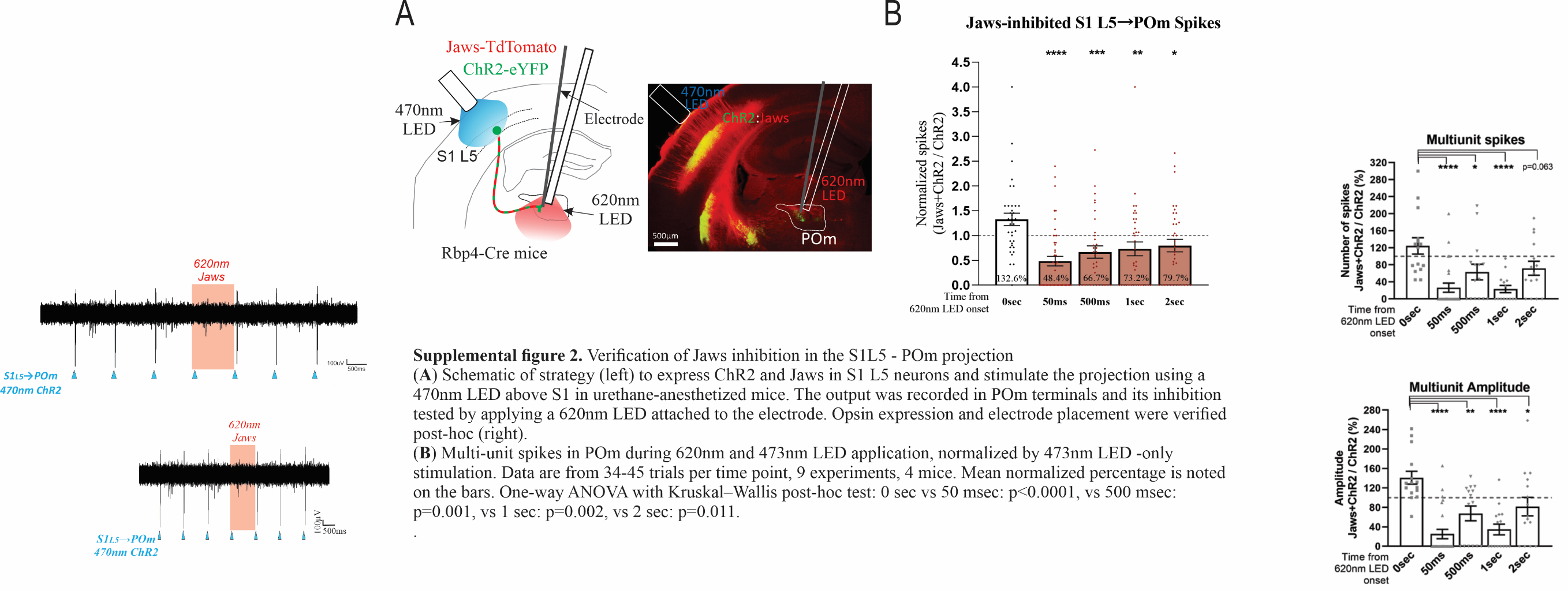

**Supplemental figure 2.** **Verification of Jaws inhibition in the S1L5 - POm projection**

(**A**) (Left) Schematic of strategy to express ChR2 and Jaws in S1 L5 neurons and stimulate the projection using a 470nm LED above S1 in urethane-anesthetized mice. The output was recorded in POm terminals and its inhibition tested by applying a 620nm LED attached to the electrode. (Right) Opsin expression and electrode placement were verified post-hoc.

(**B**) Multi-unit spikes in POm during 620nm and 473nm LED application, normalized by 473nm LED -only stimulation. Data are from 34-45 trials per time point, 9 experiments, 4 mice. Mean normalized percentage is indicated on the bars. Note that not all terminals which express ChR2 will co-express Jaws so the magnitude of suppression is likely an underestimate. One-way ANOVA with Kruskal–Wallis post-hoc test: 0 sec vs 50 msec: p<0.0001, vs 500 msec: p=0.001, vs 1 sec: p=0.002, vs 2 sec: p=0.011.

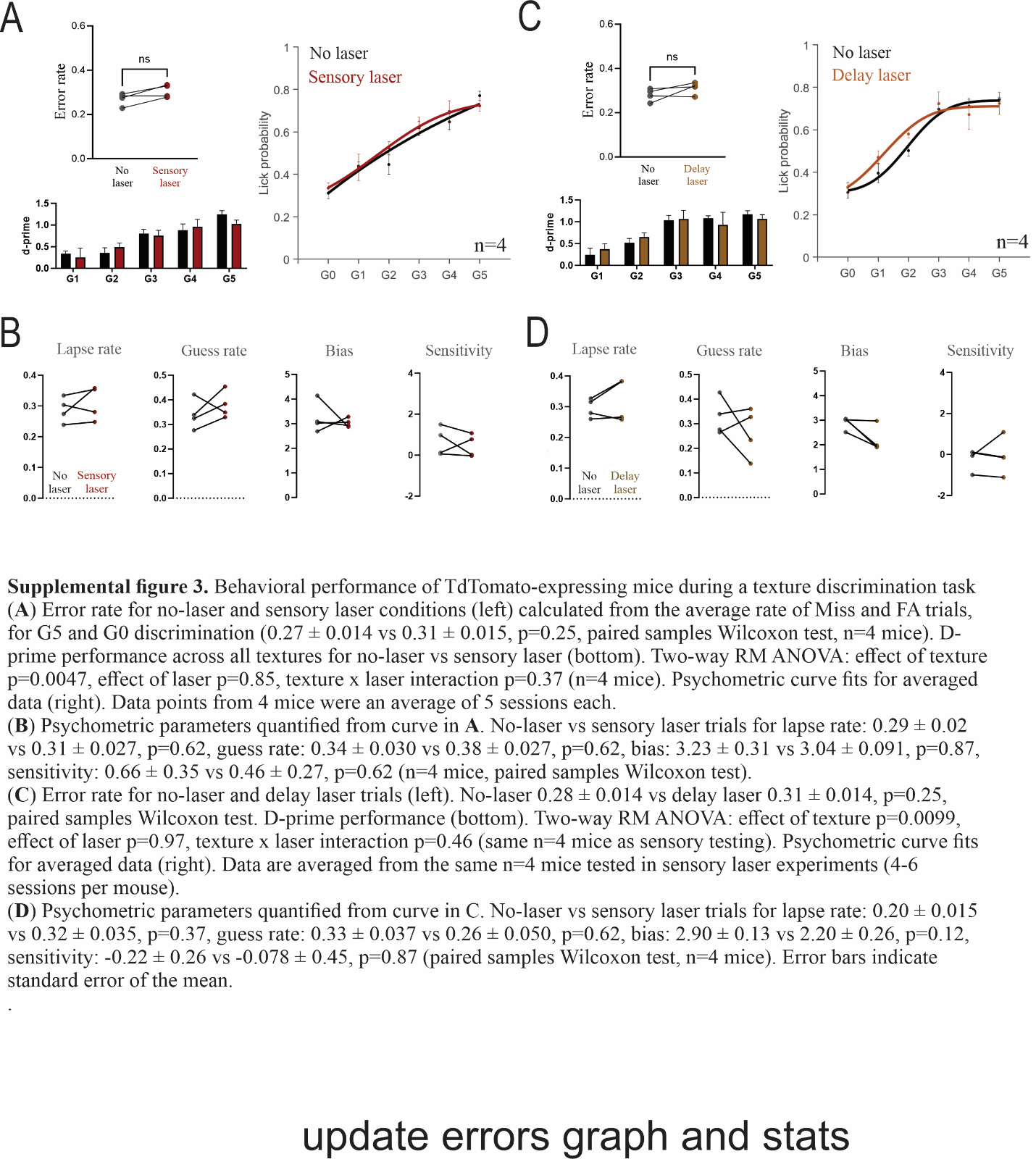

**Supplemental figure 3. Behavioral performance of TdTomato-expressing mice during a texture discrimination task**

(**A**) (Top left) Error rate for no-laser and sensory laser conditions calculated from the average rate of Miss and FA trials, for G5 and G0 discrimination (0.27 ± 0.014 vs 0.31 ± 0.015, p=0.25, paired samples Wilcoxon test, n=4 mice). (Bottom left) D-prime performance across all textures for no-laser vs sensory laser. Two-way RM ANOVA: effect of texture p=0.0047, effect of laser p=0.85, texture x laser interaction p=0.37 (n=4 mice). (Right) Psychometric curve fits for averaged data. Data points from 4 mice were an average of 5 sessions each.

(**B**) Psychometric parameters quantified from curve in A. No-laser vs sensory laser trials for lapse rate: 0.29 ± 0.02 vs 0.31 ± 0.027, p=0.62, guess rate: 0.34 ± 0.030 vs 0.38 ± 0.027, p=0.62, bias: 3.23 ± 0.31 vs 3.04 ± 0.091, p=0.87, sensitivity: 0.66 ± 0.35 vs 0.46 ± 0.27, p=0.62 (n=4 mice, paired samples Wilcoxon test).

(**C**) (Top left) Error rate for no-laser and delay laser trials. No-laser 0.28 ± 0.014 vs delay laser 0.31 ± 0.014, p=0.25, paired samples Wilcoxon test. (Bottom left) D-prime performance for no-laser vs delay laser. Two-way RM ANOVA: effect of texture p=0.0099, effect of laser p=0.97, texture x laser interaction p=0.46 (same n=4 mice as sensory testing). (Right) Psychometric curve fits for averaged data. Data are averaged from the same n=4 mice tested in sensory laser experiments (4-6 sessions per mouse).

(**D**) Psychometric parameters quantified from curve in C. No-laser vs sensory laser trials for lapse rate: 0.20 ± 0.015 vs 0.32 ± 0.035, p=0.37, guess rate: 0.33 ± 0.037 vs 0.26 ± 0.050, p=0.62, bias: 2.90 ± 0.13 vs 2.20 ± 0.26, p=0.12, sensitivity: -0.22 ± 0.26 vs -0.078 ± 0.45, p=0.87 (paired samples Wilcoxon test, n=4 mice). Error bars indicate standard error of the mean.
